# MODERATE 3D ORBITAL SHAKING MITIGATES MATERNAL SEPARATION-INDUCED NEURODEVELOPMENTAL IMPAIRMENTS IN RATS

**DOI:** 10.1101/2025.11.10.687583

**Authors:** Ulya Keskin, Melkan Kagan Kara, Onur Ozel, Cagri Caglayan Kuragi, Serel Akyol, Cansu Ozbayer, Neziha Senem-Ari, Yesim Tunc, Erkan Bayir

## Abstract

Maternal separation during early life is a well-established model for inducing neurodevelopmental impairments. Mechanical stimulation by passive movement, such as three-dimensional orbital shaking (3D-OS), has been suggested as a neuroprotective intervention in cell and tissue models, but its effects in neonatal animals remain unknown.

We investigated whether 3D-OS applied during the postnatal period could attenuate neurogenesis impairments and cognitive deficits induced by maternal separation in rats.

Eighteen newborn Sprague Dawley rats were assigned to control (C), maternal separation (MS), or maternal separation plus orbital shaking (MSOS) groups. Maternal separation was performed 3 h/day between postnatal days 2-21; MSOS additionally received 3D-OS stimulation (25 rpm, 3 h/day). Behavioural tests (Open Field, Novel Object Recognition, Morris Water Maze, Passive Avoidance), combined with RT-qPCR, Western blot, histology, and immunohistochemistry were conducted.

MSOS rats displayed improved spatial learning in the early phases of the Morris Water Maze and recovered retention memory in Passive Avoidance compared to MS. In contrast, Open Field and Novel Object Recognition tests revealed no differences, and Western blot did not detect significant protein changes. Histological analysis showed reduced neuronal loss and vacuolisation in hippocampal and cortical regions of MSOS rats. RT-qPCR demonstrated reduced hippocampal *FGF2* and *FGFR1* expression in MS, partially restored by 3D-OS, while immunohistochemistry indicated increased FGFR2 and decreased FGFR1 in MSOS.

Early-life 3D-OS administration alleviated maternal separation–induced cognitive decline and structural impairments, and enhanced hippocampal plasticity markers, providing preclinical evidence that moderate mechanical stimulation may support brain development and exert neuroprotective effects against early-life stress.

**Highlights:**

1. Early-life 3D-OS mitigated maternal separation-induced deficits.
2. Improved performance in Morris Water Maze and Passive Avoidance tasks.
3. No significant effects observed in OFT, NOR, or Western blot.
4. 3D-OS restored hippocampal FGF-2 and FGFR1 mRNA levels in stressed rats.
5. Gentle mechanical stimulation may support early brain development.

## 1. INTRODUCTION

Neurogenesis is described as novel neuron production from neural stem cells [1]. Most of the hippocampal granule neurons elongate and enhance their axons between postnatal 1st and 21st days [2]. It is also known that neurogenesis continues in adult life, decreasingly in the dentate gyrus located in the mammalian hippocampus [3,4]. Hippocampal neurogenesis is found to be associated with the amelioration of learning and memory [5]. Physical exercise has desirable impacts on the neurogenesis and, therefore, spatial learning and memory are affected positively [6,7].

It is shown that physical exercise has enhancing and improving aspects on the spatial learning and memory and increasing neuroplasticity in the preadolescent, adolescent and adult period of life, even in an environment of stress. Also, the protective effects of physical exercise on cognitive functions in neurodegenerative diseases have been noted before; besides this, recent research has accommodated physical exercise with spatial learning, especially related to the hippocampus [8,9].

Calming and getting babies to sleep via rocking a cradle is one of the methods known and frequently applied [10]. In the previous research that Inami and Koh [11] conducted, it is shown that the sleep process induced by mechanosensory stimulation has cognitive benefits on *Drosophilae*. There is research that shows vibration applications, which can be described as passive exercise, have increasing effects on neoangiogenesis and neurogenesis, and have improving effects on learning and memory [11–13]. When the method used in incubation machines is examined, turning eggs on their own axis decreases early embryo mortality and increases hatching ratios compared to the non-turned eggs. Also, it is noted that the albumen sac development is retarded, and the chorioallantoic membrane shows delayed development of 30 per cent. Besides these, the masses of the turned eggs are higher than the non-turned eggs, and the turned eggs have matured earlier [14–18]. Similarly, it is shown that 3D-OS supports neurosphere fusion and promotes the diffusion of nutrients while forming cerebral organoids in cell culture media [19]. Notably, 3D-OS provides complete conjugation and homogeneous nutrition for cells compared to uniaxial shaking [20]. When all this research is considered, mechanical stimulation applied to living cells contributes to the development process of cells in different media.

The primary role of the vestibular system (VS) is to perceive motions in spatial location and acceleration. It is shown that VS stimulation also plays a role in hippocampal cell development [21]. On the contrary, it is shown that stress, as a side note, affects VS functions both directly and indirectly [22]. Namely, stress causes impairments in the systems which affect brain development expressly or by implication. Therefore, we hypothesised that VS stimulation via 3D-OS could show ameliorating and improving effects on the neurodevelopmental impairments caused by maternal separation stress. To the best of our knowledge, no research was conducted on mechanosensory stimulation on pups during the neonatal period. Due to that, we were curious about the effects of mechanosensory stimulation on neurogenesis in babies when they cannot perform active exercise.

It is determined that there are impairments observed in neurogenesis and, hence, learning and memory in much research in which the maternal separation model is used [23–25]. Some brain chemicals play roles in neurogenesis and the neural development process. Among these chemicals, it is shown that Fibroblast Growth Factor-2 (FGF-2) shows both protective effects in the pathophysiology of neurodegenerative disorders [26] and induces proliferation [27,28]. FGF-2 induces the proliferation of progenitor cells derived from an embryonic brain or an adult hippocampus [29]. Moreover, some articles note that FGF-2 infusion into the brain increases *in vivo* neurogenesis in the dentate gyrus [30] and, also, there is an impairment observed in cortical neurogenesis in FGF-2 knock-out rats [31]. In immunohistochemical studies, it was exhibited that FGF-1 and 2 play roles in the differentiation and functions of nervous system cells, and it was seen that FGF-1 and 2 are effective in angiogenesis, smooth cell development and wound healing processes [32].

In this study, we aimed to induce an impairment in neurogenesis while inducing stress via the maternal separation method and examining the effects of 3D-OS administration on the stress-induced behavioural and molecular changes. Based on the positive effects of mechanical stimulation administered on living tissues; pups were exposed to passive exercise, which was performed by the 3D-OS method during postnatal (PN) 2-21st days before they are able to be physically active independently. It is thought that mechanical stimulation administration could primarily promote ameliorative effects on novel cell development and neurogenesis, and secondly, due to the primary concern, learning and memory are also promoted as this stimulation shows positive effects on cells [33], tissues [34] and organisms [35] in the literature. It was aimed to focus on the mechanical stimulation-induced effects of the rocking cradle-like method that is still used in our era to calm and help babies sleep, while examining and comparing the amendments in behavioural and molecular aspects.

## 2. MATERIAL AND METHODS

### 2.1. Animals and Experimental Design

All experimental procedures were performed in accordance with the Guide for the Care and Use of Laboratory Animals [36] and were approved by the Kütahya Health Sciences University Animal Experiments Local Ethics Committee (protocol no: 2024.12.02, approval date: December 2, 2024).

In this study, 18 newborn Sprague Dawley rats were used. Both sexes were balanced across groups (3 males and 3 females per group) and were not separated, as the animals were in the prepubertal stage when the influence of gonadal hormones is minimal until the puberty onset, postnatal 40 to 60th days [37–40]. Rats were housed under controlled conditions (20-22 °C, 12 h light/12 h dark cycle) with free access to standard laboratory chow and water *ad libitum*. Lights were on at 07:00 and off at 19:00. All behavioural tests were conducted between 10:00 and 15:00 under stable environmental conditions.

Rats were randomly separated into three groups:

1. Control Group (C) (n=6): No administration performed.
2. Maternal Separation Group (MS) (n=6): Exposed to stress by getting separated for 3 hours from their mothers between PN 2nd-21st days.
3. Maternal Separation Stress + Orbital Shaking Group (MSOS) (n=6): Additionally, to the same stress protocol, exposed to 3D-OS administration.

#### Maternal Separation Protocol

Between PN 2-21st days, every day for 21 days, the mother rat has been moved to another cage, without touching or handling the pups and kept waiting for 3 hours in an environment where the temperature is controlled by an air conditioner between 32 ± 0.5 °C [41–43]. By this protocol, stress was established in pups. Thus, impairment in the neurogenesis was cellularly determined and anxiety and learning, which came up as a result of this impairment, were evaluated behaviourally.

#### Shaking Protocol

For the same number of days in the same environment mentioned above, pups were put onto a 3D orbital shaker (LCD Digital 3D Shaker Dlab SK-D3309-Pro) in their cage, which were fixed on the shaker table, for 3 hours and shaken with a 9° tilt angle and 25 rpm frequency of rotation to create a rocking cradle-like effect [19,44,45]. Pups were not stabilised, or no contact was made by researchers in order to take precautions against mother rats rejecting pups. Protective or ameliorating effects of this procedure on impaired neurogenesis were evaluated.

#### Sacrification Protocol

Before the sacrification procedure, rats were anaesthetised using a combination of ketamine (90 mg/kg) and xylazine (10 mg/kg). After the induction of anaesthesia, rats were sacrificed by the exsanguination method. Brain samples were collected after the sacrifice, and the brains were cut between the hemispheres into two equal pieces. One piece was put into the 10% neutral buffered formalin solution for histological evaluation, and the other piece was kept in -80°C for molecular tests.

### 2.2. Behavioural Tests

All behavioural tests were conducted from the 35^th^ day of the experiment onwards. One day before each test, the rats were placed in the experimental apparatus for five minutes to complete the acclimatisation process. The experiments in which images are recorded are done with a DMK 22AUC03 monochrome camera, and the software produced by the academicians and the students of Kutahya Health Sciences University, Faculty of Engineering and Natural Sciences.

#### 2.2.1. Open Field (OF) Test

Each rat was placed at the exact centre of an OF apparatus, which was a square prism made of wood, 60 x 60 cm at the base and 35 cm high, with an open top, and was observed for 300 seconds [46]. Anxiety levels of the rats were assessed by determination of locations (centre or peripheral areas) that rats prefer to occupy in the apparatus.

#### 2.2.2. Novel Object Recognition (NOR) Test

The basis of the mechanism of this test depends on rats becoming interested and getting more curious in the novel object rather than the familiar one. In the first trial of the experiment, the rats were allowed to spend 5 minutes with two of the same objects. After the rat underwent the first trial, one of the objects was changed to a new one. When the change was made, the rat was placed back into the apparatus, and the time spent with the novel object was evaluated. The two objects used in the experiment were similar in height, volume but different from each other in colour and shape. Exploring behaviour was counted when the rat was 2-3 cm maximum away from one of the objects, touched one of them, sniffed or did vibrissae sweeping. Without exploring behaviour, the time the rats were occupying a place near or above one of the objects was not counted and evaluated [47]. In the protocol, the total time spent with both of the objects and the time with the first object that the rat had interacted with were assessed.

The discrimination index (*DI*) is used to evaluate NORT results and formulated as *DI* = (*T*_novel_ - *T*_familiar_) / (*T*_novel_ + *T*_familiar_), where *T* denotes exploration time [25].

#### 2.2.3. Morris Water Maze (MWM) Test

MWM tests were done in a tank with a diameter of 130 cm and a depth of 60 cm, filled with water at a temperature of 24±2 °C, with a 10 cm diameter escape platform placed 1 cm below the water level. The position of the platform was not changed during the whole experiment, and cues were placed outside the water tank at a distance and height visible to the rats.

MWM was used to assess hippocampal learning, and the experiment that lasted for 5 days was ended for the rats which could not find the platform within 120 seconds in each trial [48,49]. Rats were placed into the water maze from four different directions as north, south, west and east, in each trial; hence, rats were precluded from memorising the escape track, and were required to utilise their spatial learning abilities by observing and learning the cues. In the first trial, the rats which were not able to find the platform in 120 seconds were gently pointed in the right direction and were aided in finding the platform by a stick. Each time they found the platform, the rats were allowed to stay there for 15 seconds to comprehend the cues spatially. For 5 days, learning was evaluated by the behavioural performance that the subjects exhibited. During these evaluations, the time it took for the rats to find the platform (latency time) was recorded, and the percentage of time spent in the target quadrant was calculated using the following formula: (time spent in the target quadrant/time spent in the other quadrants) × 100.

#### 2.2.4. Passive Avoidance (PA) Test

PA experimental apparatus (Passive Avoidance Step-Through Cat. No. 40550) consists of two chambers, one being dark, and in addition, is set up to apply an electrical shock, and a light one wherein no shock is given. The floor of the PA apparatus is made from stainless steel grating, and there is a passage door between the chambers that can be opened. In the first trial, 60 seconds were given as a waiting time for the rats to explore the environment, and the door was arranged to open at the 60th second. As rats passed to the dark chamber, 0.4 milliamperes of electrical shock was applied for 3 seconds by the grating. Transition times from the light chamber to the dark chamber (latency) were evaluated after 1 hour (first day, second trial) for short-term memory assessment and after 24 hours (second day, third trial) for medium-long-term memory assessment. To evaluate the latency, the cut-off value was determined as 300 seconds [50–52].

### 2.3. Cellular Experiments

#### 2.3.1. Real-Time Polymerase Chain Reaction (RT-qPCR)

RT-qPCR protocol consists of RNA isolation, cDNA synthesis and qPCR stages for target gene expression (*FGF2*, *FGFR1* and *FGFR2*). Total RNA was isolated from the hippocampal samples that were preserved at -80°C by using RNA isolation kits provided by the producer. After isolation, RNA concentrations were measured using Nanodrop (Allsheng Instruments, China). The cDNAs were synthesised by using a cDNA synthesis kit recommended by the producer and a thermal cycler (T100 Thermal Cycler, Bio-Rad, USA), from RNA samples which were standardised according to their concentrations. cDNA synthesis was checked using Nanodrop (Allsheng Instruments, China).

RT-qPCR mix was prepared using β*-actin* primers as a housekeeping gene, forward and reverse primers for target genes (*FGF2*, *FGFR1*, *FGFR2*). The reaction was prepared for each sample as follows: 5 μl SYBR Green Mix, 6.5 μl molecular biological grade water, 1.5 μl cDNA, 1 μl Forward Primer, and 1 μl Reverse Primer. The prepared reaction mixtures were loaded into a 96-well plate, and the plate surface was covered with a transparent film. The RT-qPCR reaction was amplified with an initial denaturation of 10 minutes at 95°C, followed by 35 cycles of 95°C for 15 seconds and 60°C for 10 seconds (StepOne-Plus Real-Time PCR System). After RT-qPCR amplification, ΔCт and 2^-ΔΔCт methods were used to assess target gene expression levels. Specific primer sequences used in this study are given in Table 1.

**Table 1.**
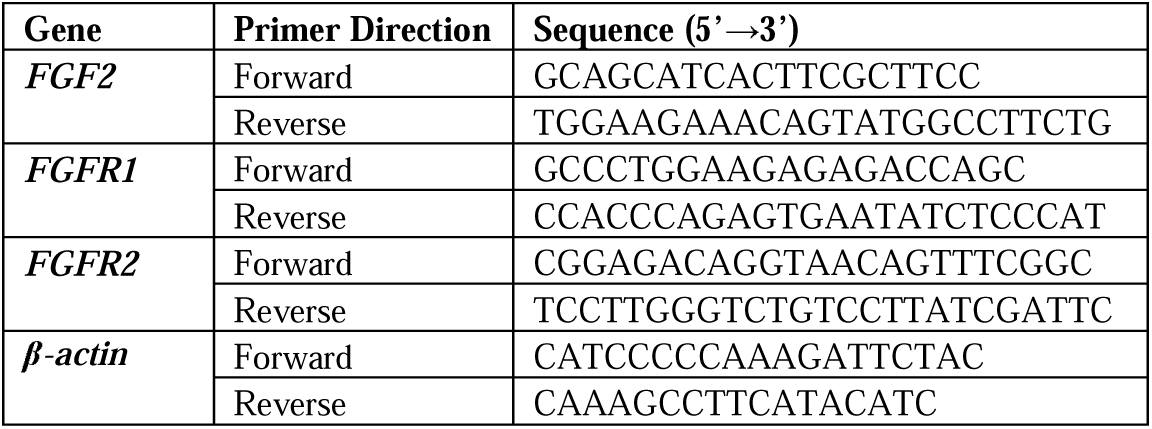
Primer sequences used for quantitative RT-qPCR analysis.

#### 2.3.2. Western-Blot Analysis

Determination of FGFR1 and FGFR2 protein levels using Western Blot consists of the following steps: protein extraction, total protein assay, SDS-PAGE (Sodium Dodecyl Sulphate - Polyacrylamide Gel Electrophoresis), transfer to the membrane, primary and secondary antibody incubation and chemiluminescence imaging. Protein extracts were prepared from the hippocampal samples by a commercial homogenization buffer (xTractor™ Buffer; Takara Bio, Mountain View, CA) per kit protocol and using TissueLyser (Qiagen, Valencia, CA, USA) homogeniser. Total protein level of protein lysates was determined spectrophotometrically at 595 nm by using protein standards prepared with Bradford protein assay and bovine serum albumin (BioRad Laboratories). After samples were standardised (50 μg) accordingly to their total protein levels, it was passed to the SDS-PAGE stage. Following the preparation of resolving and stacking gels (FastCast Acrylamide Solutions, Bio-Rad, USA), standardised samples were mixed with loading buffer, denatured by heat, loaded to SDS-PAGE and electrophoresis was performed (Mini-Protean® Tetra system, Bio-Rad, USA). After electrophoresis, a sandwich was formed with gel, filter paper and PVDF (Polyvinylidene Fluoride) membrane, and transfer was performed using the Trans-Blot Turbo Transfer System (Bio-Rad).

After blocking the membranes with a blocking solution to determine target protein levels, primary antibodies for FGFR1 and FGFR2 proteins and HRP-conjugated secondary antibodies were incubated and then imaged using a Chemiluminescence imaging system (ChemiDoc MP imaging system, Bio-Rad, USA) with luminol reagent (Clarity Max Western ECL Substrate, Bio-Rad). The images were analysed using ImageLab software and calculated as raw/vol, then normalised to β*-actin*.

#### 2.3.3. Histology and Immunohistochemistry

Tissues were fixed for 48 hours in 10% neutral buffered formalin for microscopic examination. Following the fixation, tissues were washed under flowing tap water for 24 hours. After the tissues were passed through increasing degrees of alcohol solutions (80%, 96%, 98%, 100%) to become dehydrated, they were cleaned in xylene until transparent, and then they were embedded in paraffin blocks. 5 μm slices, obtained by using a fully automatic microtome device, were placed onto a glass slide and stained using the Haematoxylin-Eosin (HE) staining method to assess cortical and hippocampal architecture. Additionally, the samples obtained were submerged two times in xylene for a duration of 30 minutes each, subsequent to being stored in an oven maintained at 60 °C. Then, rehydration was achieved with alcohol series ranging from 95 % to 60 %, which were then incubated in distilled water for 5 minutes and held at room temperature for 15 minutes in a 0.5% trypsin solution, limited with Pap Pen. To inhibit endogenous tissue peroxidase, a 3% HLOL solution was applied for 5 minutes. The sections were washed three times for 5 minutes each with phosphate buffer solution (PBS) and then treated with blocking solution for 1 hour. After the blocking solution was removed from the tissue, the sections were incubated with primary antibodies overnight. The next day, the sections were washed three times with buffer solution and stained with secondary antibodies for 30 minutes. The sections were washed three times with buffer solution for 5 minutes each and stained with DAB (3,3′-diaminobenzidine) for 5 minutes to determine the visibility of the immunohistochemical reaction. After staining with Mayer’s Haematoxylin, the sections were washed with distilled water for 10 minutes and sealed with sealing medium. The resulting preparations were evaluated under a microscope (Zeiss Axiovision, Germany) by two histologists in a double-blind manner.

### 2.4. Statistical Analysis

Sample size was calculated as 18 animals in total (α = 0.05, power = 80%, effect size *f* = 0.9), with three groups designed for the study. For descriptive statistics, arithmetic mean, standard deviation, minimum and maximum values were reported for quantitative data, and frequency and percentage distributions were provided for qualitative data. Normality of the distributions was assessed using the Shapiro-Wilk test. For normally distributed data, one-way ANOVA was applied, and for non-normally distributed data, the Kruskal-Wallis test was used. When overall group differences were significant, pairwise comparisons were performed: Tukey’s post hoc test after ANOVA, and Bonferroni-corrected Mann-Whitney *U* test after Kruskal-Wallis. In figures, data are presented as mean ± standard error of the mean (SEM) for normally distributed variables and as median (IQR) for non-parametric data. Statistical significance was set as *p* < 0.05.

## 3. RESULTS

The data that support the findings of this study are available from the corresponding author upon reasonable request.

### 3.1. Behavioural Experiments

#### 3.1.1. Open Field

One-way ANOVA revealed no significant difference in OFT scores among the groups, *F*(2,15) = 0.98, *p* = 0.399. Mean (±SD) OFT values were 13.21 ± 8.90 for the C, 8.20 ± 7.41 for the MS, and 8.47 ± 3.46 for the MSOS.

#### 3.1.2. Novel Object Recognition

One-way ANOVA revealed no significant difference in NORT (familiar object exploration) values among the groups, *F*(2,14) = 0.57, *p* = 0.578. Mean (±SD) NORT scores were 60.54 ± 68.44 for the C, 45.52 ± 22.49 for the MS, and 88.39 ± 98.19 for the MSOS.

One-way ANOVA revealed no significant difference in DI scores among the groups, *F*(2,14) = 0.52, *p* = 0.608. Mean (±SD) DI values were -0.47 ± 0.72 for the C, -0.16 ± 0.35 for the MS, and -0.21 ± 0.52 for the MSOS.

One-way ANOVA revealed that latency time on the first day of MWM experiments was significantly longer in MS compared to C (*F*(2,15) = 16.74, *p* < 0.001) and in MSOS compared to C (*p* = 0.004), whereas no difference was observed between MS and MSOS (*p* = 0.238; Figure 1A). On the third day, the escape latency in the MS was significantly longer compared to C (*F*(2,15) = 4.98, *p* = 0.022). Although the difference between MS and MSOS did not reach statistical significance (*p* = 0.057), it showed a trend toward longer latency in the MS, while no difference was observed between C and MSOS (*p* = 0.925; Figure 1A), indicating only a non-significant trend toward recovery with 3D-OS.

**Figure 1.**
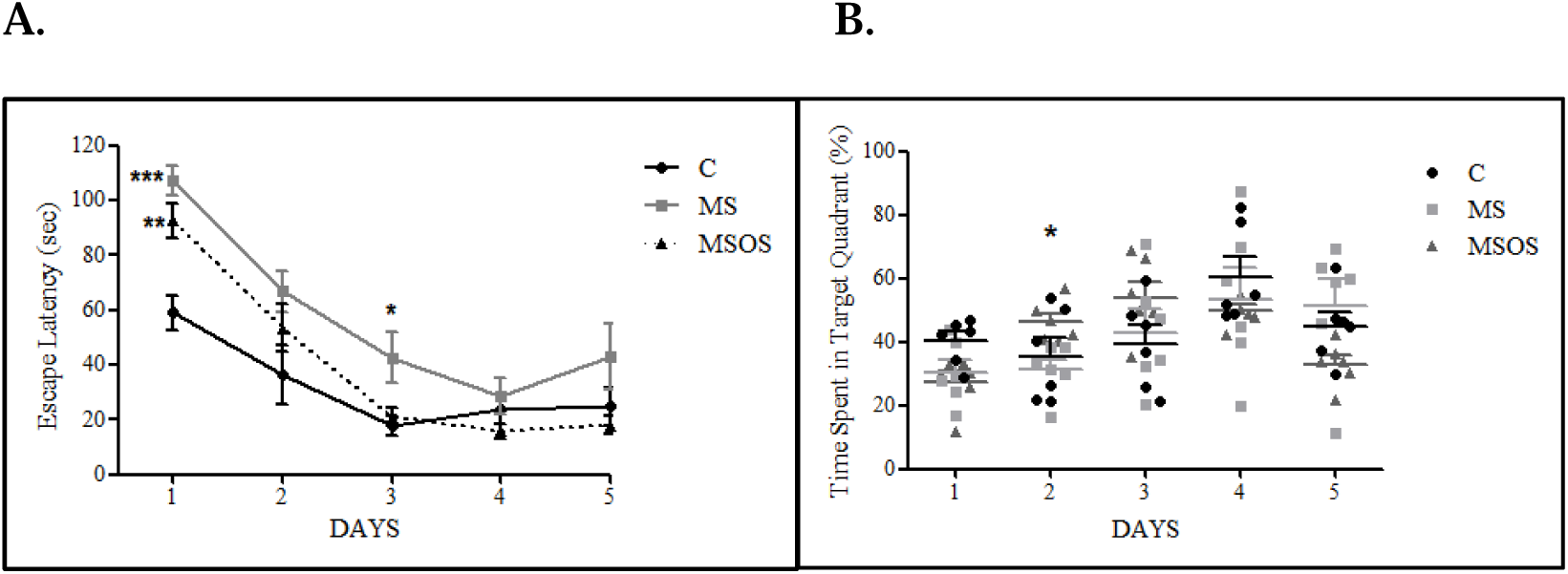
(A) Escape latency in the Morris Water Maze (MWM) test over 5 consecutive days (\**p* < 0.05, \*\**p* < 0.01, \*\*\**p* < 0.001 vs C). Data are presented as mean ± SEM. (B) Percentage of time spent in the target quadrant over 5 days (\**p* < 0.05 vs C; ±*p* < 0.05 vs MS). Data are presented as individual values with the median (horizontal line) and interquartile range (Q1-Q3). C: Control; MS: Maternal Separation; MSOS: Maternal Separation with Orbital Shaking (*n* = 6 per group).

The percentage of time spent in the target quadrant was calculated by dividing the time spent by the rats in the quadrant where the platform was located by the total swimming time. Median (Q1-Q3) values for the time spent in the target quadrant are presented in Table 2. On the second day, the Kruskal-Wallis test showed a significant group difference (*H*(2) = 6.33, *p* = 0.042). Post hoc pairwise comparisons using the Mann-Whitney *U* test revealed that MSOS spent a significantly greater proportion of time in the target quadrant compared to MS (*U* = 0.00, *Z* = −2.88, *p* = 0.004), while no significant differences were found between C and MS (*p* = 0.631) or between C and MSOS (*p* = 0.262; Figure 1B).

#### 3.1.3. Passive Avoidance

In the second trial (T2), no statistical significant difference was observed among the experimental groups in step-through latency (*H*(2) = 5.23, *p* = 0.073; Figure 2), although the MS showed a tendency toward shorter latency compared with the C, indicating a mild impairment in avoidance learning that did not reach statistical significance.

**Figure 2.**
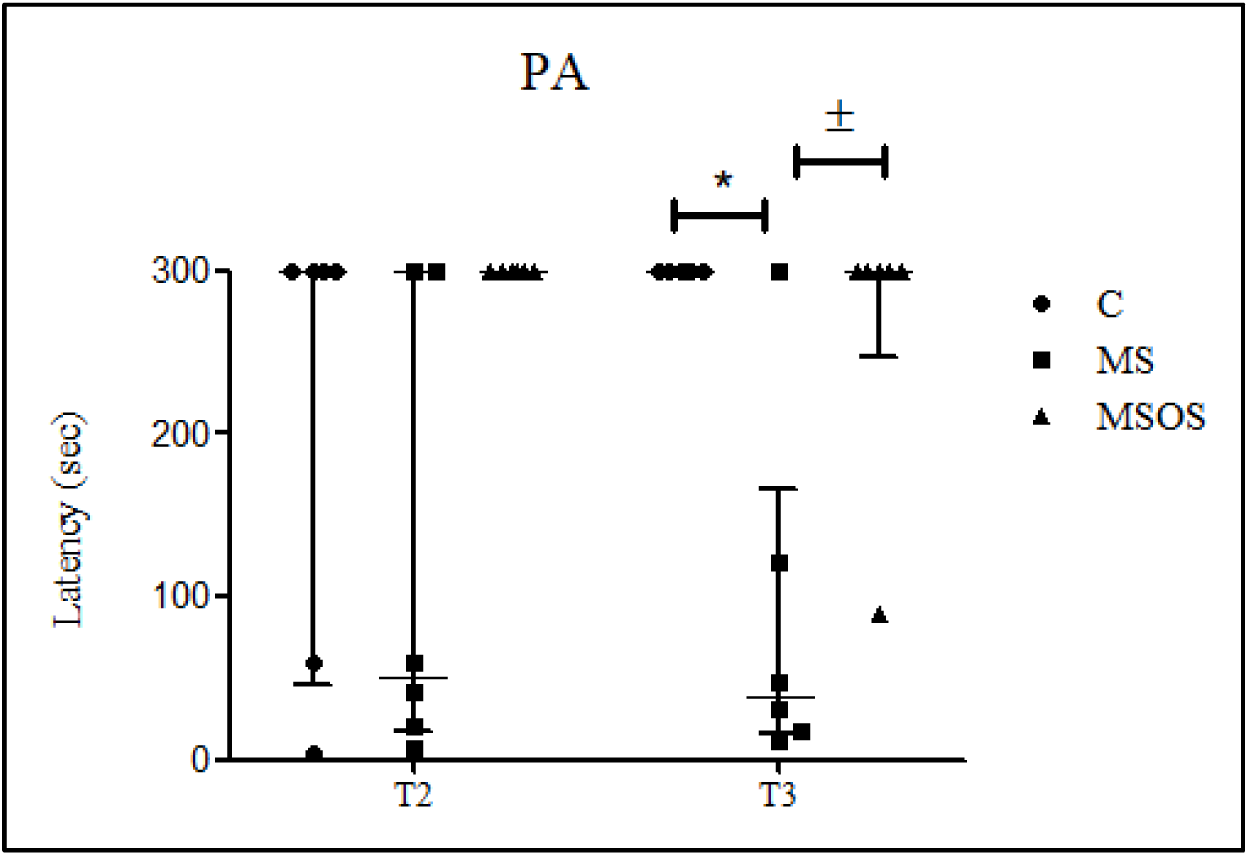
Latency in the Passive Avoidance (PA) Test. Data are presented as individual values with the median (horizontal line) and interquartile range (IQR, whiskers) (*n* = 6 per group). In the third trial (T3), MS showed significantly shorter latency compared with C (\**p* < 0.05) and MSOS (±*p* < 0.05). C: Control; MS: Maternal Separation; MSOS: Maternal Separation with Orbital Shaking (*n* = 6 per group).

In the third trial (T3), however, a significant group difference was detected in step-through latency (*H*(2) = 10.34, *p* = 0.006; Figure 2). As illustrated in Figure 2, T3 step-through latencies were markedly shorter in MS and restored by 3D-OS. Post hoc pairwise comparisons using the Mann-Whitney *U* test revealed that the MS exhibited materially shorter latencies than both the C (*U* = 3.00, *p* = 0.007) and the MSOS (*U* = 4.50, *p* = 0.021). No significant difference was observed between the C and MSOS (*U* = 15.00, *p* = 0.317). These findings indicate that maternal separation substantially impaired retention memory, whereas 3D-OS effectively reversed this deficit.

### 3.2. Cellular Experiments

#### 3.2.1. Real-Time Polymerase Chain Reaction (RT-qPCR)

Relative mRNA expression levels of *FGF2*, *FGFR1*, and *FGFR2* were analyzed using the Kruskal-Wallis test followed by Bonferroni-corrected Mann-Whitney U tests for pairwise comparisons (*n* = 6 per group). Significant group differences were observed for *FGF2* (*H*(2) = 8.43, *p* = 0.015) and *FGFR1* (*H*(2) = 10.19, *p* = 0.006), whereas *FGFR2* expression did not differ significantly among groups (*H*(2) = 1.91, *p* = 0.39; Figure 3A–C). Post hoc analyses revealed that the MS exhibited significantly lower *FGF2* and *FGFR1* mRNA levels compared with both the C (*p* = 0.010 for *FGF2*; *p* = 0.010 for *FGFR1*) and the MSOS (*p* = 0.016 for *FGF2*; *p* = 0.004 for *FGFR1*), while no differences were found between the C and MSOS (*p* = 0.873 for both genes). These results indicate that maternal separation significantly reduced *FGF2* and *FGFR1* transcription, whereas early-life orbital shaking stimulation restored their expression to near-control levels, with no significant changes observed in *FGFR2*.

**Figure 3.**
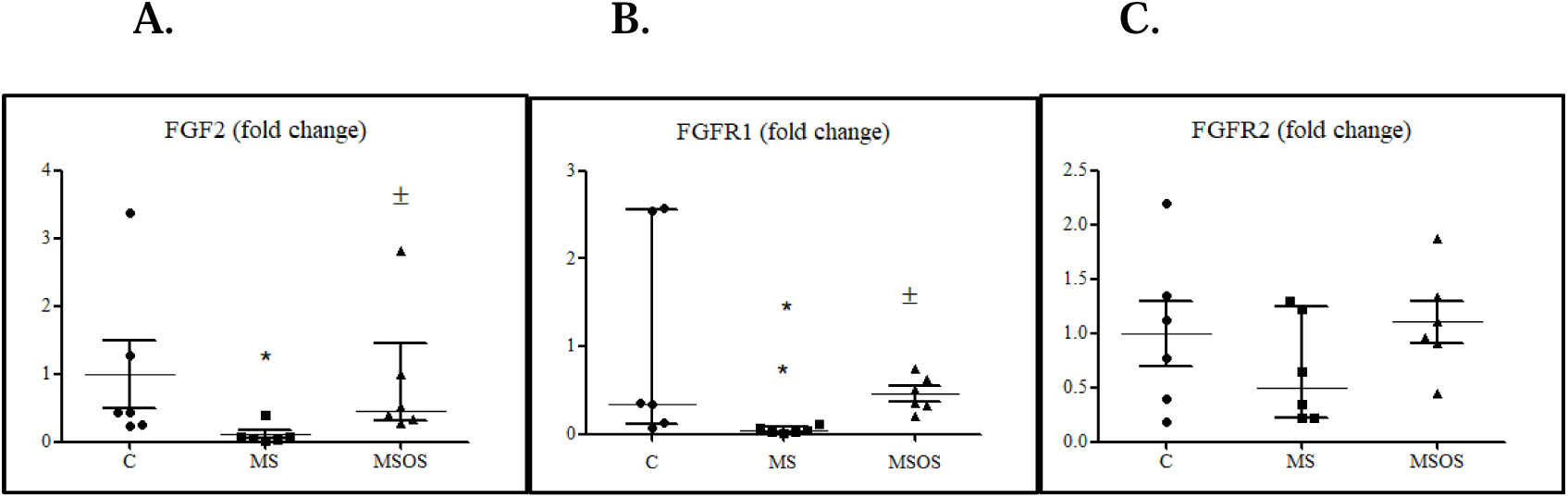
RT-qPCR analysis results of *FGF2* (A), *FGFR1* (B), and *FGFR2* (C) mRNA expression in whole-brain homogenates. Data were normalised to β*-actin* using the 2^−ΔΔCt method and are presented as individual values with the median (horizontal line) and interquartile range (IQR, whiskers) (*n* = 6 per group). C: Control; MS: Maternal Separation; MSOS: Maternal Separation with Orbital Shaking (*n* = 6 per group). Significant differences: \**p* < 0.05 vs C; ±*p* < 0.05 vs MS.

#### 3.2.2. Western-Blot Analysis

Western blot analyses were performed on a randomly selected subset of animals (*n* = 3 per group). Protein expression levels of FGFR1 and FGFR2 in whole-brain homogenates were compared among groups using the Kruskal-Wallis test. No statistically significant differences were found for either FGFR1 (*H*(2) = 1.07, *p* = 0.587) or FGFR2 (*H*(2) = 2.49, *p* = 0.288; Figure 4A-B).

**Figure 4.**
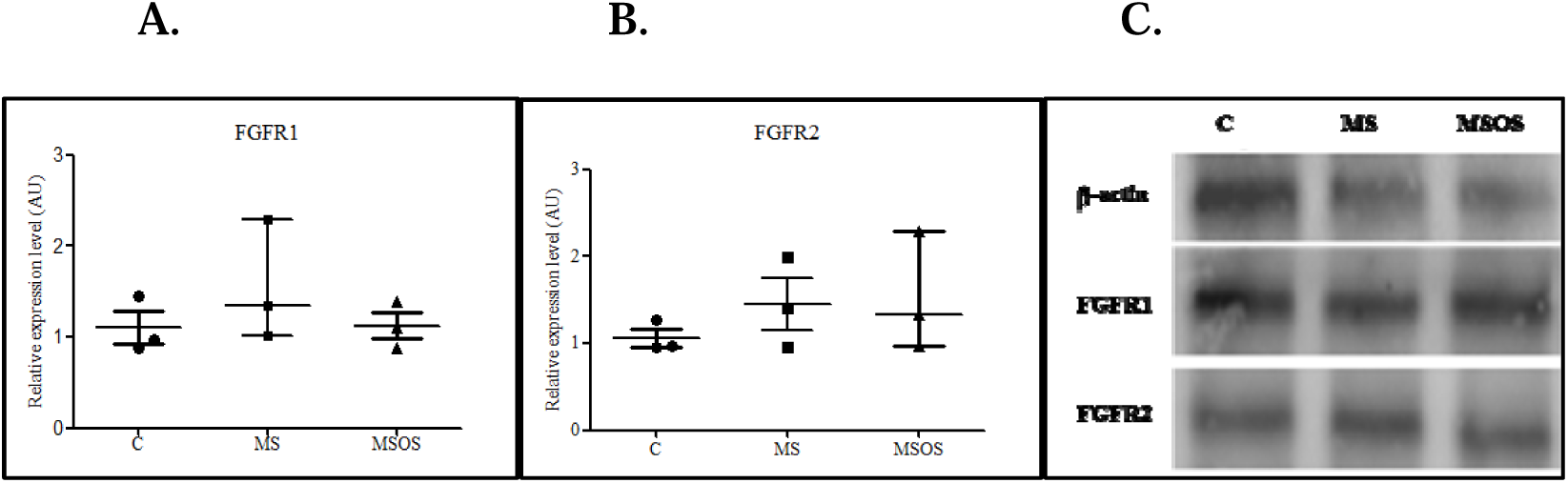
(A) Densitometric quantification of FGFR1, (B) FGFR2 protein levels in whole-brain homogenates, and (C) representative Western blot bands showing FGFR1 and FGFR2 expression normalised to β*-actin*. Data are presented as individual values with the median (horizontal line) and interquartile range (IQR, whiskers) (*n* = 3 per group). C: Control; MS: Maternal Separation; MSOS: Maternal Separation with Orbital Shaking.

Although these changes did not reach statistical significance, FGFR1 levels appeared approximately 40% higher in the MS compared with C, and FGFR2 levels were modestly elevated in both MS and MSOS. These findings suggest a non-significant upward trend in FGFR signalling following maternal separation and partial normalisation with 3D-OS.

#### 3.2.3. Histology

##### 3.2.3.1. Cortex Assessment

Normal histology was observed in samples belonging to C. In MS, neurons were surrounded by vacuoles, and apoptotic glial cells were shown. Cells which are surrounded by vacuoles were fewer and sporadic in MSOS compared to MS.

##### 3.2.3.2. Hippocampus Assessment

In the CA1 region of MS and MSOS, numbers of neuron layers were decreased compared to C. Also, in the CA3 region of MS, the number of neurons was decreased compared to MSOS and C (Figure 5).

**Figure 5.**
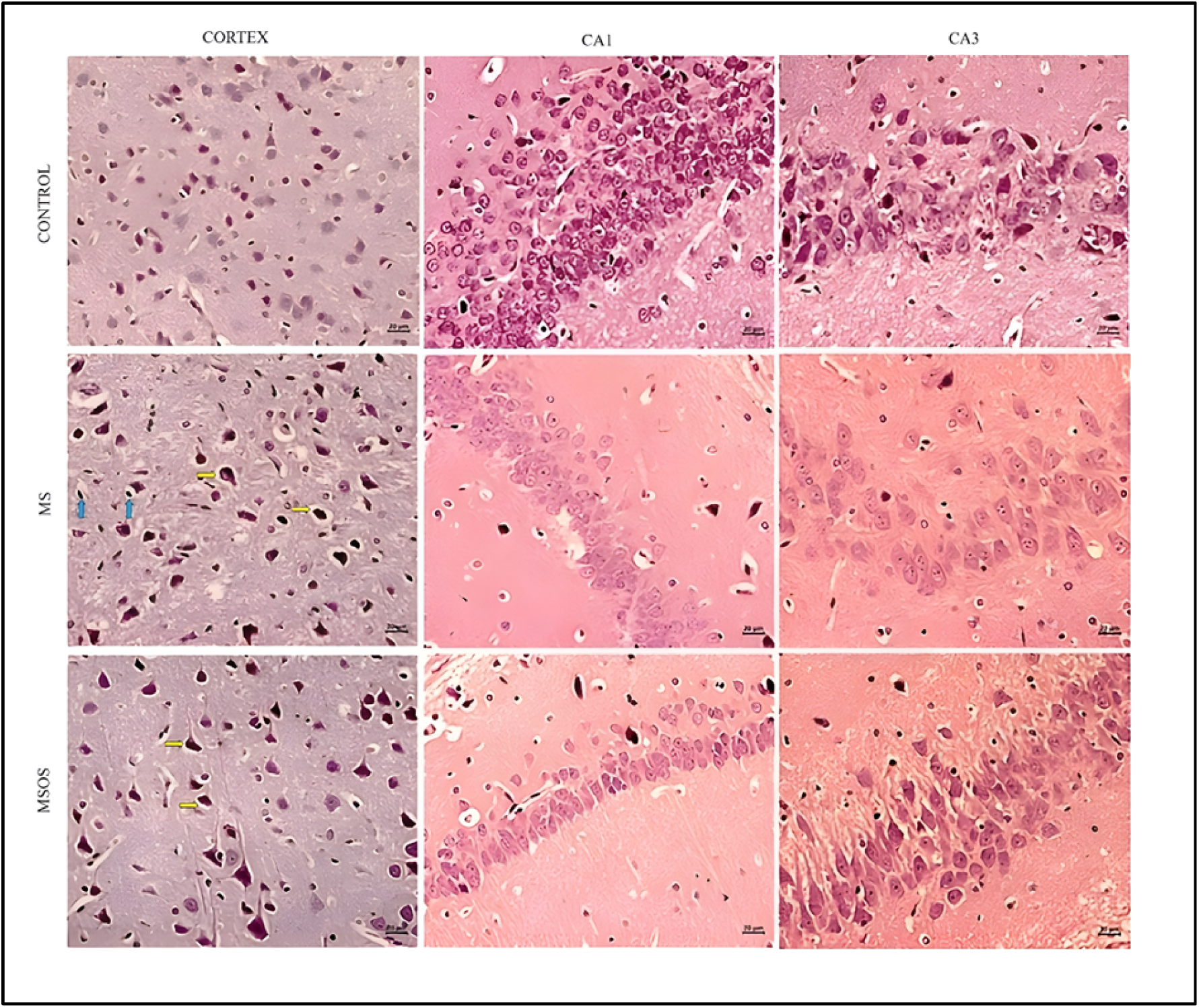
Histological sections from cortical and hippocampal areas. Columns represent the cortex (left), hippocampal CA1 (middle), and CA3 (right) regions. Rows represent the C (Control), MS (Maternal Separation), and MSOS (Maternal Separation with Orbital Shaking) groups (*n* = 6 per group). Yellow arrows indicate neurons surrounded by vacuoles, and blue arrows indicate apoptotic glial cells. Scale bar: 20 µm.

#### 3.2.4. Immunohistochemistry

Quantitative analysis of immunostaining revealed significant group differences in FGFR1 and FGFR2 expression within hippocampal sections (*n* = 6 per group).

For FGFR1, one-way ANOVA indicated a robust group effect (*F*(2,15) = 194.46, *p* < 0.001; Figure 6A). Tukey’s post hoc comparisons showed markedly reduced FGFR1 immunoreactivity in both MS (*p* < 0.001) and MSOS (*p* < 0.001) groups relative to C, with the reduction most pronounced in the MSOS.

**Figure 6.**
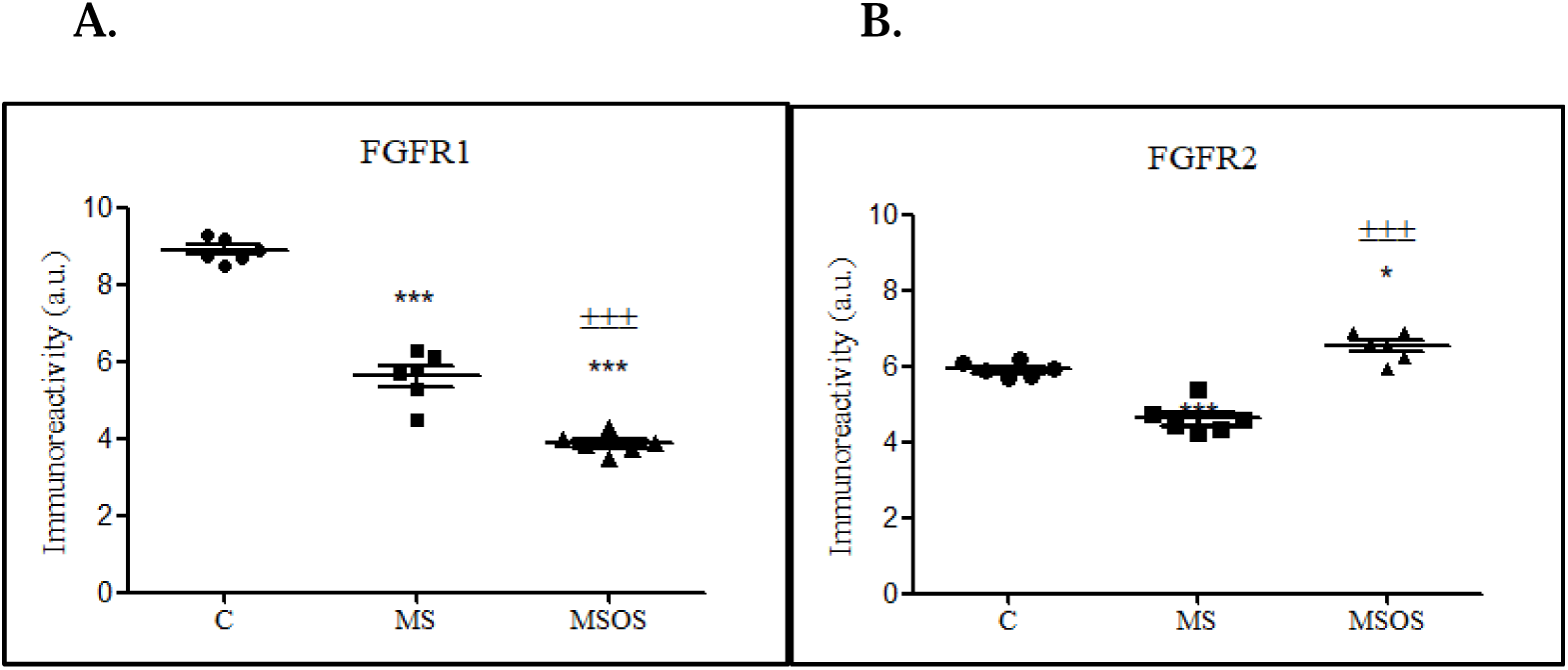
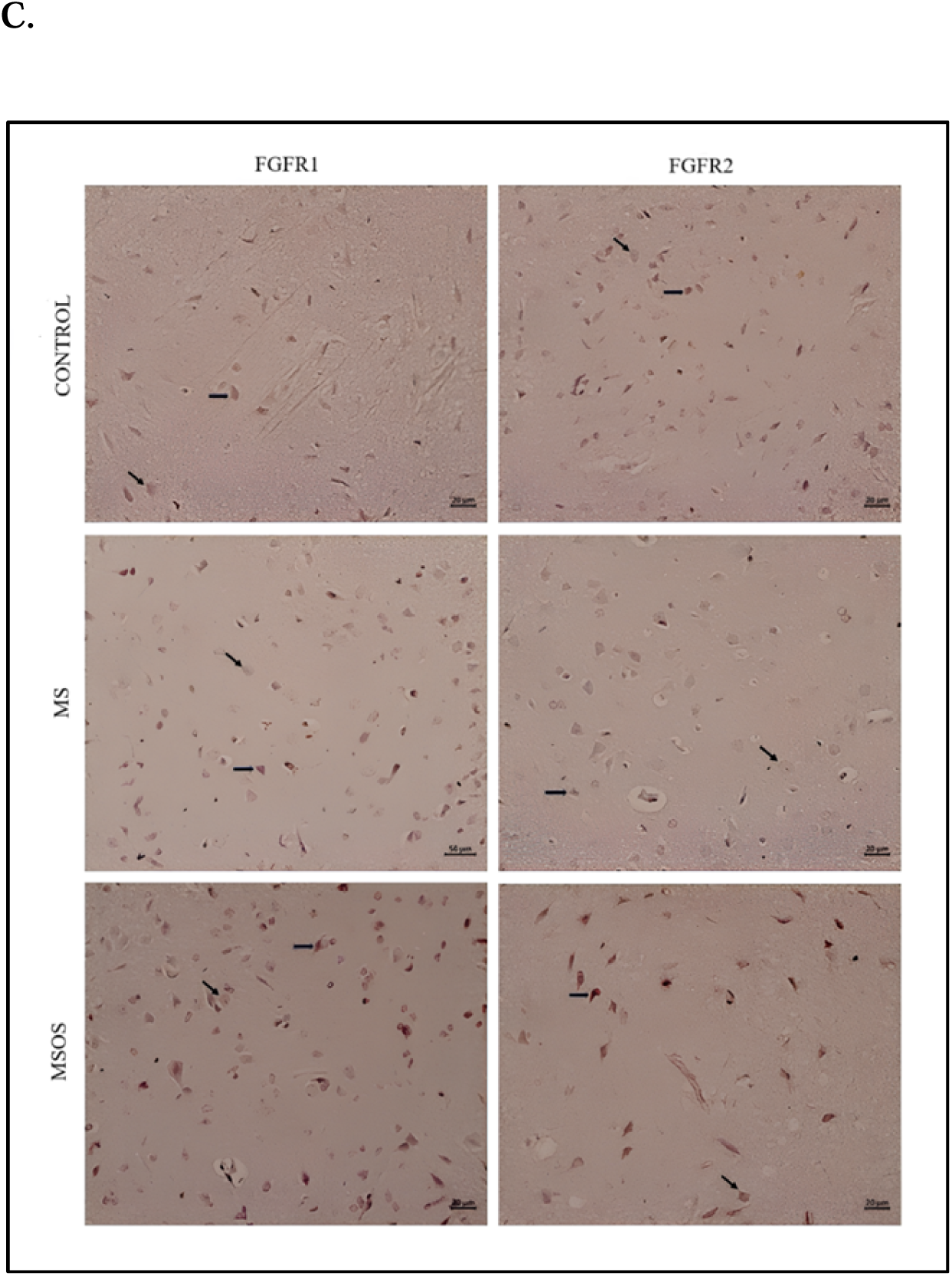
Quantitative and qualitative analysis of FGFR1 and FGFR2 immunoreactivity in the hippocampus. (A) FGFR1 and (B) FGFR2 immunoreactivity levels are shown as individual values (mean ± SEM, *n* = 6). (C) Representative hippocampal CA images of FGFR1 (left) and FGFR2 (right) immunostaining in C (Control), MS (Maternal Separation), and MSOS (Maternal Separation with Orbital Shaking) groups. Black arrows indicate positively stained neurons. Scale bar: 20 µm. Significant differences: \**p* < 0.05 vs C; \*\*\**p* < 0.001 vs C; ±±±*p* < 0.001 vs MS.

In contrast, FGFR2 expression also differed significantly among groups (*F*(2,15) = 49.68, *p* < 0.001; Figure 6B). The MS exhibited a strong reduction in FGFR2 immunopositivity compared with the C (*p* < 0.001) and MSOS (*p* < 0.001), whereas FGFR2 staining intensity in the MSOS exceeded that of C (*p* = 0.019).

These findings suggest that maternal separation markedly suppressed both FGFR1 and FGFR2 immunoreactivity, while early-life orbital shaking stimulation counteracted these reductions in a receptor-specific manner, leading to partial recovery of FGFR1 and a trend toward increased FGFR2 expression, as also illustrated by representative hippocampal immunostaining images (Figure 6C).

## 4. DISCUSSION

This study aims to explore the effects of 3D-OS administration on the developmental neurogenesis and cognitive impairments depending on maternal separation stress, behaviourally, molecularly and histologically. Rats participating in the study were not separated according to their genders since they were in the prepubertal stage when gonadal hormonal impact is minimal and can be neglected [39,40]. The results obtained from this study show that 3D-OS administration may alleviate cognitive impairments caused by maternal separation stress and positively affect hippocampal plasticity.

In various research, it has been proven that maternal separation stress causes impairing effects on hippocampal functions, and this detrimentally affects the learning and memory processes [53,54]. The decrement observed in the behavioural performance in MWM and PA tests was similarly marked in our study [55]. Consistent improvements in both MWM and PA suggest that 3D-OS enhances hippocampal-amygdalar integration underlying spatial and avoidance memory. In MWM, escape latency increased in MS; deficits showed partial recovery with MSOS approaching C over days. As shown in Figure 1A, escape latencies were elevated in MS and showed only a partial, time-dependent recovery in MSOS relative to C. While target-quadrant preference improved significantly on day 2 compared with MS, escape-latency differences between MS and MSOS did not reach significance. This pattern is consistent with the target-quadrant preference shown in Figure 1B. This recovery supports the hypothesis that 3D-OS administration may enhance hippocampal plasticity via the vestibular system [56,57].

The increase in target quadrant orientation observed in the MSOS suggests that 3D-OS partially corrects stress-induced spatial learning impairments (Figure 1B). The findings indicate that 3D-OS administration enhances cognitive performance, as evidenced by the observation that MSOS spent a longer duration in the target quadrant on the second day in comparison to MS (*p* = 0.004) and showed a tendency toward longer target quadrant duration on the third day in relation to C (*p* = 0.078). By contrast, the disappearance of statistical differences on the fourth and fifth days can be explained by the fact that the learning levels of all groups approached each other over time. This condition can be associated with the ceiling effect or plateau formation in the learning curve, where further improvement is limited due to maximal task acquisition [58]. Similarly, when escape latency data were compared (Figure 1A), on the 1st to 3rd days, the slope of the learning curve was steeper and latency was higher. The fact that the difference between groups disappears on the fourth day, presumably due to the ceiling effect, also attracts attention. In this context, the tendency to swim in a target-oriented manner, which was observed in the early period seems to substantiate the alleviating potential of 3D-OS administration on the neurobiological effects caused by stress.

It is known that maternal separation stress suppresses the neurogenesis in the dentate gyrus, therefore, affects the learning and memory processes negatively [23]. The evidence showed that the mechanical stimulation may promote the neuronal plasticity via mechanosensitive ion channels such as Piezo1/2 [59]. Effects of mechanosensory stimulation established by 3D-OS in the initial period of life can be interrelated with its regulatory role on hippocampal neurogenesis. Given the critical role of FGF2 in progenitor proliferation and stress resilience, its recovery by 3D-OS indicates a possible upstream modulation of the FGFR1 signalling axis.

It has been observed that 3D-OS administration has improving effects on learning and memory tests, but creates no significant difference in the tests that focus on anxiety [60] and episodic-like declarative memory [61], such as OF and NOR. This suggests that 3D-OS may affect spatial learning and short- to medium-term memory processes, which are more closely related to hippocampal functions. The absence of significant differences in OF and NOR tests suggests that 3D-OS mainly influences hippocampus-dependent spatial learning (MWM) and amygdala-dependent inhibition/avoidance memory (PA) modulation, as indicated by improvements in MWM and PA tasks, respectively. However, 3D-OS does not affect general locomotory activity (OFT) or episodic-like declarative memory processes (NOR).

The behavioural recovery seen in MWM and PA tests is also promoted at the cellular and molecular levels. In molecular analysis, it was observed that *FGFR1* mRNA expression is reduced in MS; however, it was partly improved by 3D-OS administration. It is known that FGFR1 signalling plays a crucial role in neuronal proliferation and synaptogenesis processes [62]. Within this context, although *FGFR1* transcription appeared restored in MSOS, this recovery did not consistently translate into protein expression. This suggests that 3D-OS could exert its effects primarily at the transcriptional or post-transcriptional level, while translational regulatory mechanisms may limit protein availability. Therewithal, it suggests that in Western blot analysis, the FGFR1 protein levels could not reach a significant difference due to increases at the transcriptional stage that were not reflected in translation, and that translational suppression mechanisms might be active together with post-transcriptional remodelling [63].

FGFR1 protein expression was found to be lower in both MS and MSOS compared to C in immunohistochemical analysis, hence it exhibits a discrepancy with RT-qPCR results. These kinds of discrepancies are frequently reported in the literature, and can be interpreted and explained by transcriptional and post-translational regulation mechanisms such as translational suppression, differences in mRNA structure stability, microRNA-mediated regulations and cell type-specific expression differences [63–65]. This finding indicates the importance of post-translational regulations under stress conditions. To explore these mechanisms, more studies need to be conducted on this topic.

In our study, an increase in FGFR2 protein levels was detected only by immunohistochemical staining in the MSOS, whereas qPCR and Western blot analyses did not confirm this finding. This discrepancy may reflect cell type-specific or localized protein expression changes, or methodological sensitivity, rather than a uniform compensatory mechanism. Notably, the divergent pattern of decreased FGFR1 and increased FGFR2 (only in IHC) may suggest a receptor-level compensatory balance; however, this remains speculative and requires further validation in future studies [62,64].

The mechanical stimulation established by 3D-OS may trigger the cellular response by being transmitted to the hippocampus via vestibular pathways. This process may regulate neurotrophic factors and, thus, support the recovery. Mechanical inputs in the brain tissue are translated to biochemical signals by mechanotransduction; the mechanically sensitive channels and receptors fire this signalling and trigger downstream responses [66]. These mechanical signals may be affecting processes such as synaptic plasticity and neurogenesis [67]. In this perspective, the increment seen in FGFR2 can be assessed as associated with the establishment of compensatory responses and the involvement of secondary signalling pathways. Also, mechanosensitive channels (i.g, Piezo channels) are likely to mediate these processes; the Piezo family were described as primary transducing sensors which translate the mechanical stimulation into electrochemical signals and play a role in many cell types in the central nervous system [68]. Despite the fact that we have not investigated these mechanisms in our study, the explanations provided here should be seen as a mechanistic hypothesis that correlates with our data and has to be tested in the future, particularly via the study designs which directly explore mechanical sensitivity and alterations in the receptor level.

Histopathological examinations also substantiate these results at the structural level. Vacuolisation, neuronal loss and morphological impairments were observed in hippocampal and cortical regions of MS; besides, these findings were determined to be explicitly reduced in MSOS. The neuron number was particularly preserved in the hippocampal CA3 region, which points to the fact that mechanical stimulation protects developing brain tissue against stress. This finding supports the neuroprotective effects of 3D-OS at the microscopic level [62,64]. 25 rpm of 3D-OS velocity used in this study has not been directly tested on animal behaviour models. However, the decision for this speed was made considering the studies conducted on cell culture and organoid development [33]. The orbital shaking protocols, which Watanabe et al. [20] used in their organoid creation studies, both provide gentle stimulation and preserve the cellular integrity. In a similar manner, regularly turning avian eggs affects embryonic development in a positive way. The 25 rpm of velocity we chose was assessed as a gentle stimulator for the vestibular system, but a mechanical stimulation level which does not create stress. Comparison of different rpms is going to be made in the future, which can prove whether this effect is frequency-dependent or not. It was shown that, as a similar mechanical stimulation method, the whole body vibration (WBV) supports cognitive processes as neurogenesis, angiogenesis and synaptic plasticity in animal models [13,69]. These findings prove that mechanical stimulation can be protective and enhancing in the nervous system. Thus, provides indirect evidence that supports the potential benefits of 3D-OS.

The genuine part of this study in the literature is that the 3D-OS was used for the first time in an animal model and assessed the effects with behavioural, molecular and histological results. In the current literature there is indirect evidence which shows, shaking is used widely for putting babies to sleep [10], orbital shaking increases cellular fusion and differentiation [19], turning optimises the embryo development in the avian incubations [15]; besides these, this study was the first to show that effects of this kind of mechanical stimulation on neurodevelopmental process.

Some limitations are also present in our study. Firstly, the mechanisms underlying the effects of mechanical stimulation, such as vestibular system activation, neurotransmitter levels or stress hormones (e.g. corticosterone), were not directly measured. In addition, FGFR levels were only shown in mRNA and immunohistochemical levels, but cellular proliferation or neurogenesis markers were not evaluated by biochemical methods. This has prevented the underlying neuronal changes in observed behavioural effects from being explained in more detail. Notwithstanding all these limitations, our study presents the potential of 3D-OS as a novel environmental and behavioural intervention method. In future studies, the comparison between different frequencies, times and administration periods of 3D-OS; differentiation of effects depending on age and gender, measuring stress hormones and use of neurogenesis biomarkers will be enlightening. Moreover, in translational research, how this kind of mechanical stimulation corresponds with vestibular stimulation or sensorimotor therapy modalities in humans should be examined. These findings provide a preclinical framework for neonatal vestibular or gentle sensory stimulation approaches to promote neurodevelopment in at-risk infants.

Although the molecular mechanisms remain to be fully elucidated, the convergence of behavioural, molecular, and histological improvements indicates a consistent pattern that strongly needs further investigation of the 3D-OS effects.

## 5. CONCLUSION

This study suggests that 3D-OS applied during the neonatal period may mitigate stress-induced impairments in learning, memory, and support neurodevelopmental resillience. These beneficial effects were associated with an increase in *FGFR1* expression at the transcript level without consistent confirmation at the protein level, reduced neuronal injury in hippocampal subfields, and improved performance in hippocampus-dependent behavioural tasks (MWM, PA). As a novel paradigm of gentle mechanical stimulation, 3D-OS may enhance neuroplasticity and act as a protective intervention against early-life stress. Future studies combining behavioural, molecular, and electrophysiological approaches are needed to clarify the mechanistic basis of these effects and to determine whether this approach holds translational potential for innovative interventions in neurodevelopmental disorders in the future.

## 8. ACKNOWLEDGEMENTS

We would like to thank the employees of the Experimental Animal Unit, Faculty of Medicine, Kutahya Health Sciences University for their technical support and to Susan Savaş for her language support.

## 9. DECLARATION OF GENERATIVE-AI OR AI-ASSISTED TECHNOLOGIES USED IN THE STUDY

During the process of writing this article ChatGPT, OpenAI was used merely for grammatical corrections and refining the sentences for fluency. AI has not been involved in any processes of acquiring, creating, interpreting, visualising or expressing the data. All authors have revised the final version of the paper and verified the manuscript to ensure its accuracy and factuality. All authors take the full responsibility for the content of this publication.

